## Supplementary Figures 1-19 for "Deep screening of proximal and distal splicing-regulatory elements in a native sequence context"

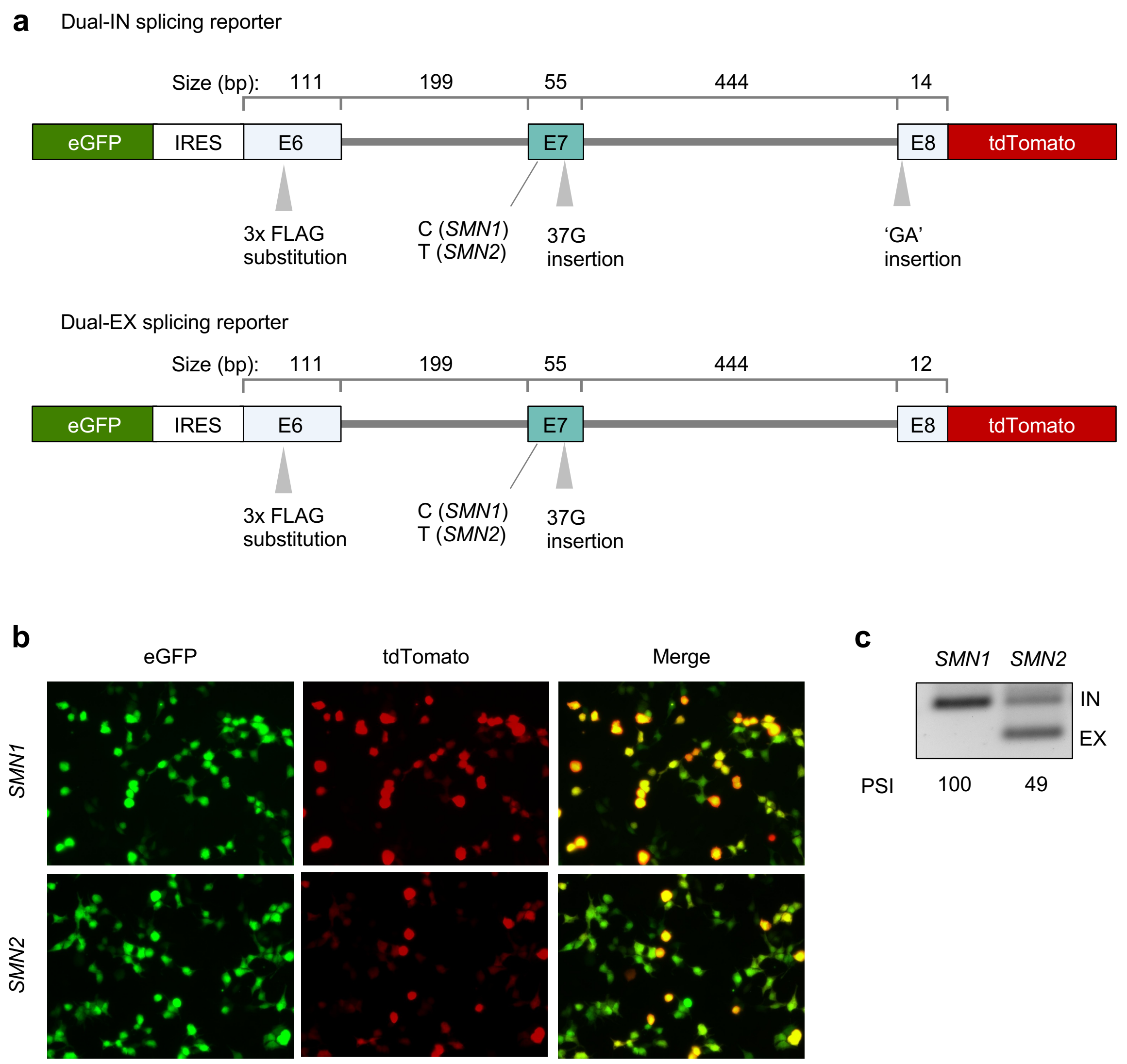


Supplementary Figure 1

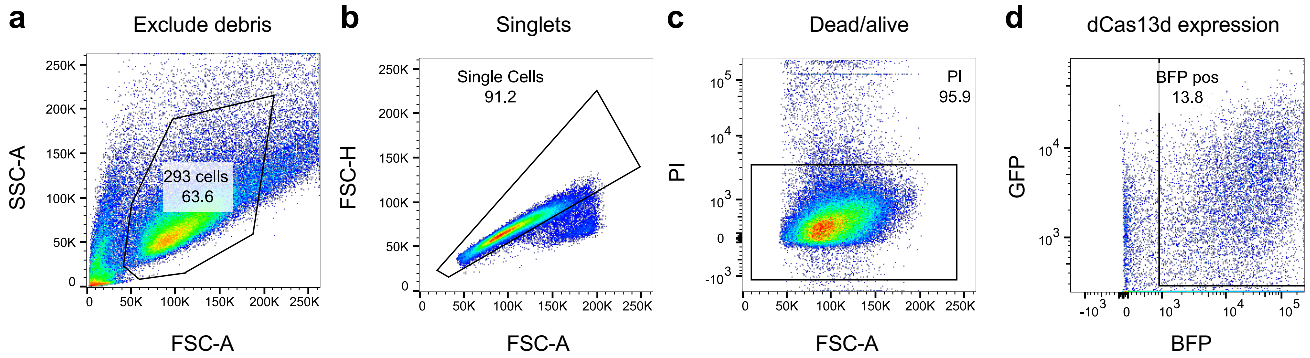


Supplementary Figure 2

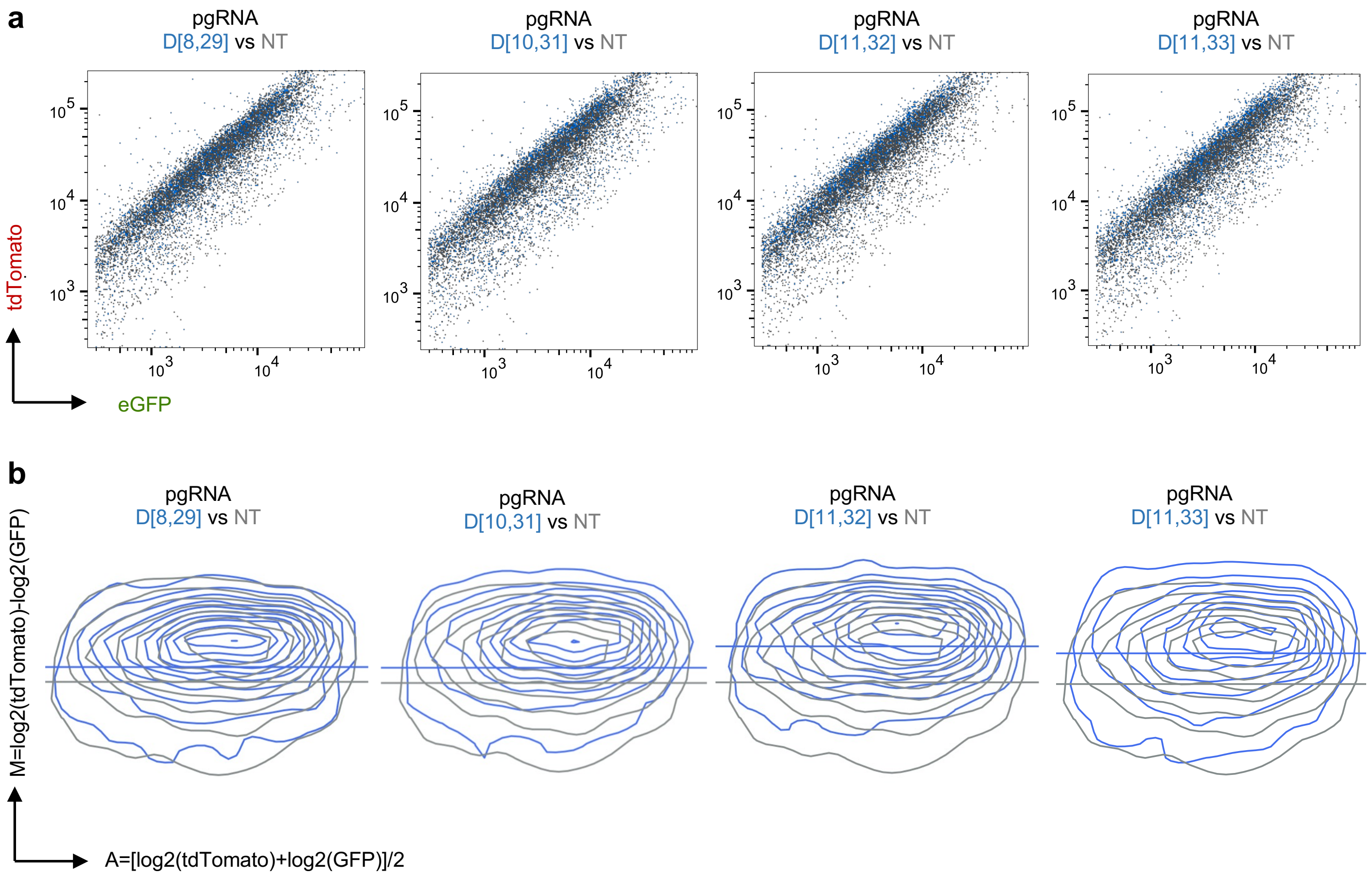


Supplementary Figure 3

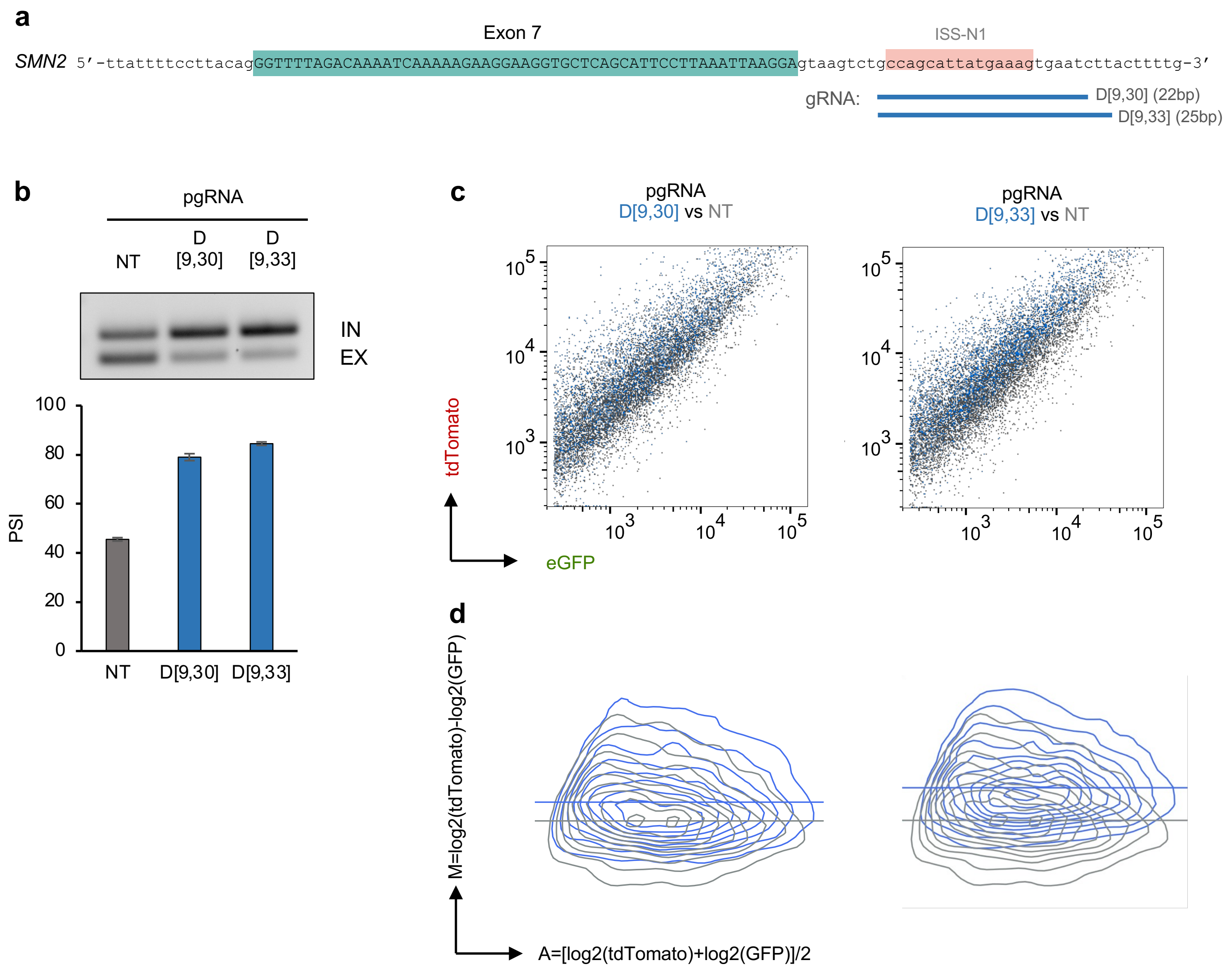


Supplementary Figure 4

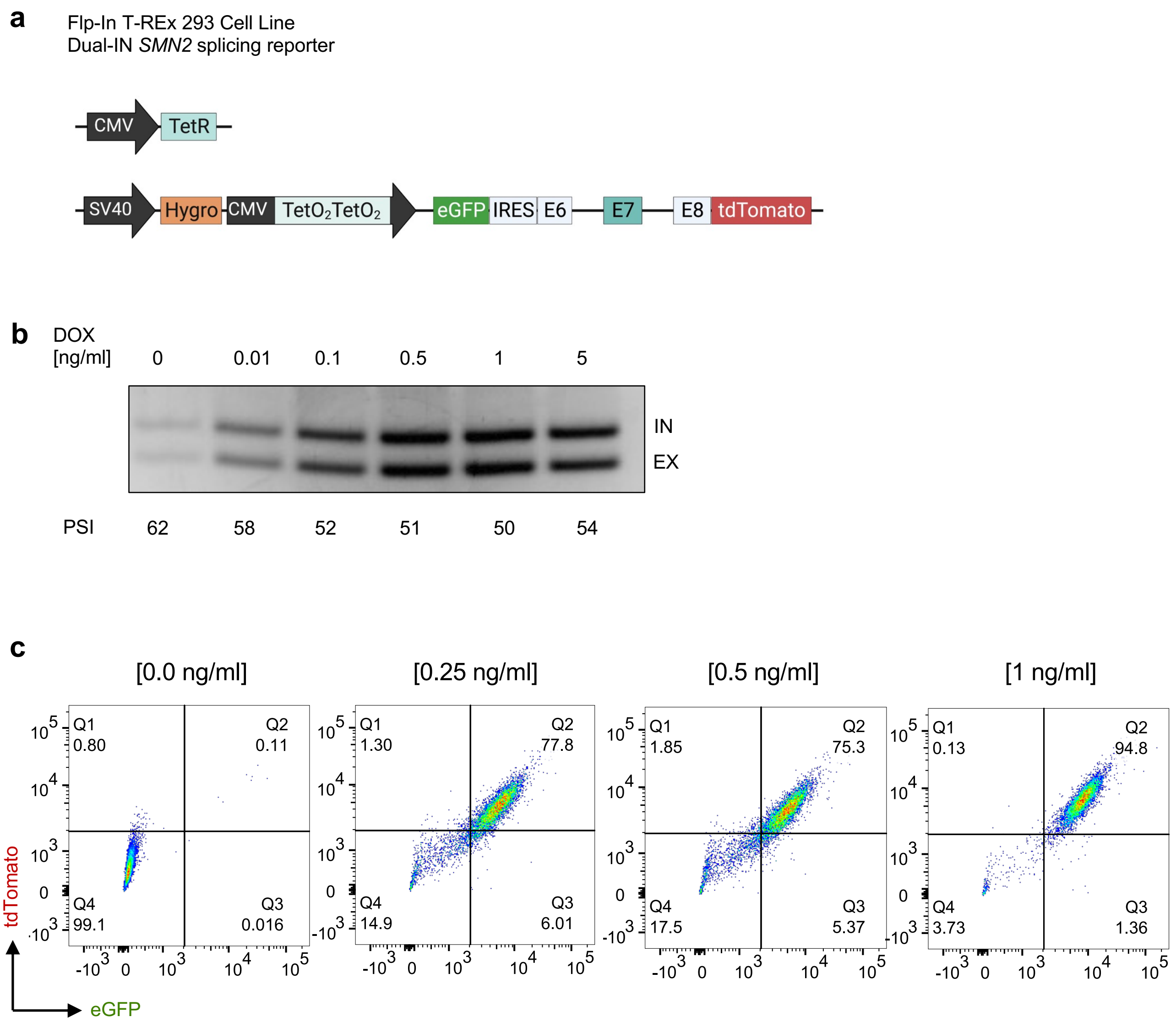


Supplementary Figure 5

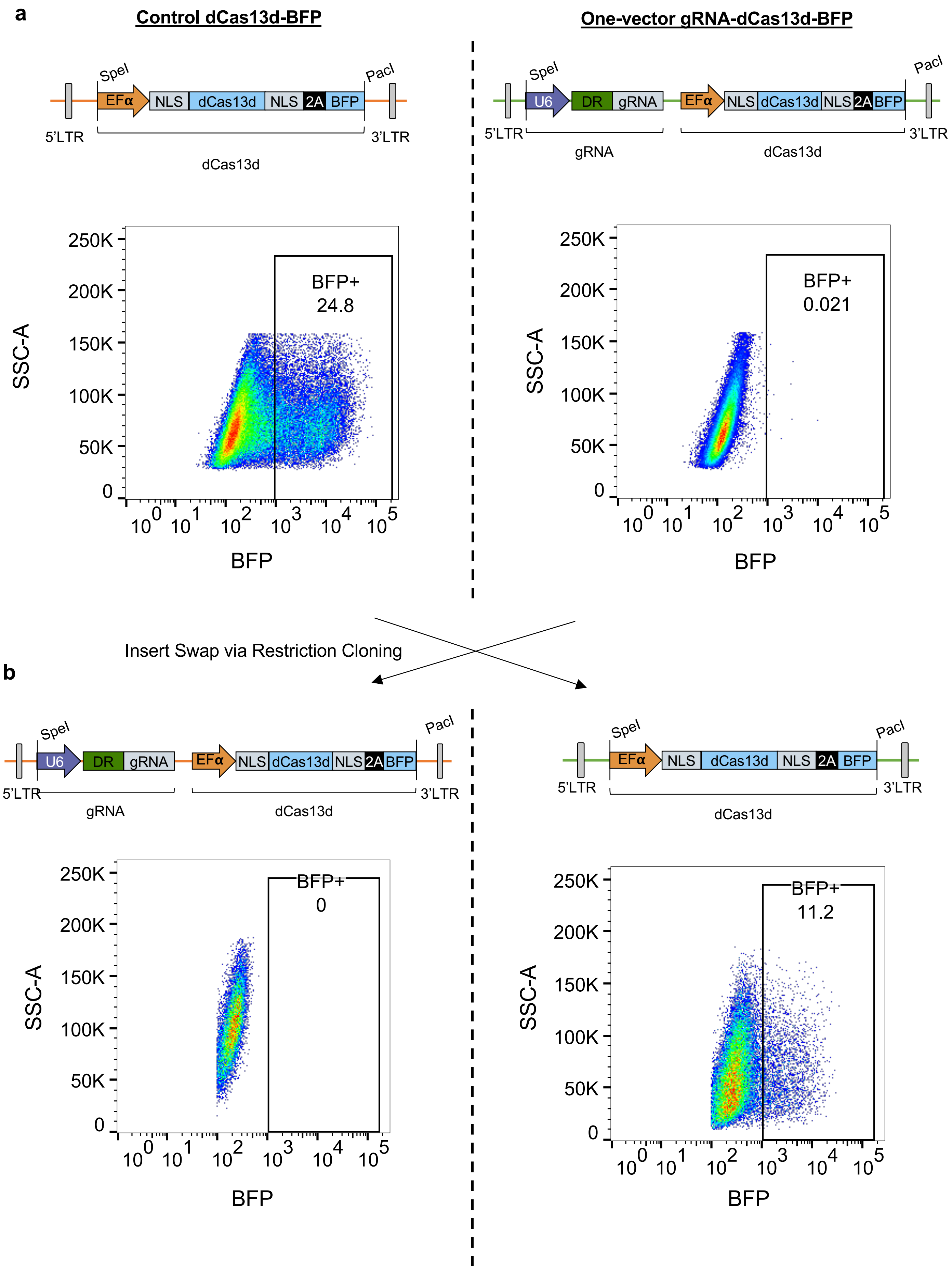


Supplementary Figure 6

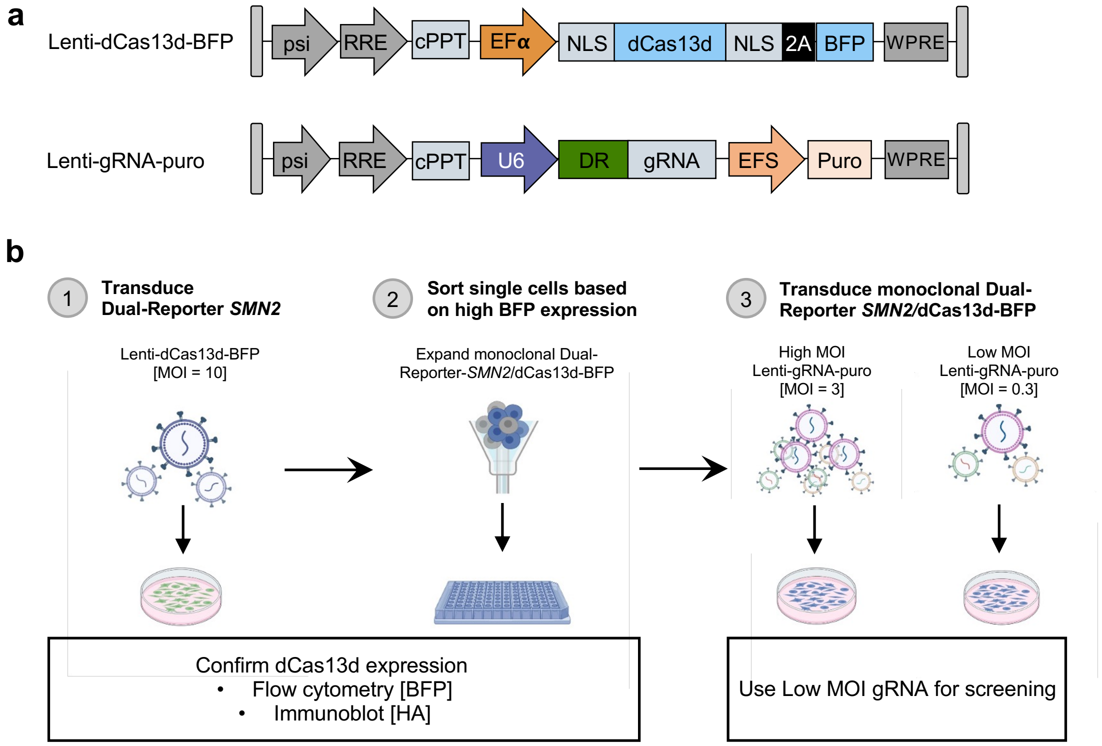


Supplementary Figure 7

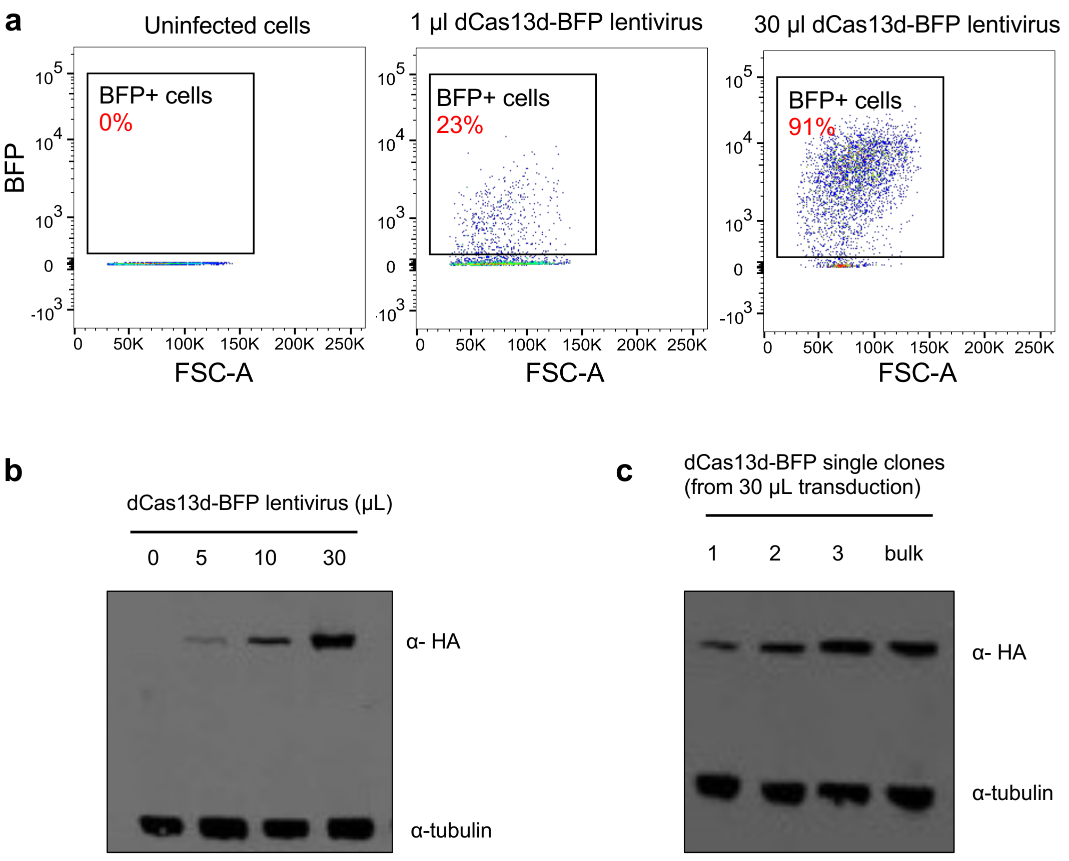


Supplementary Figure 8

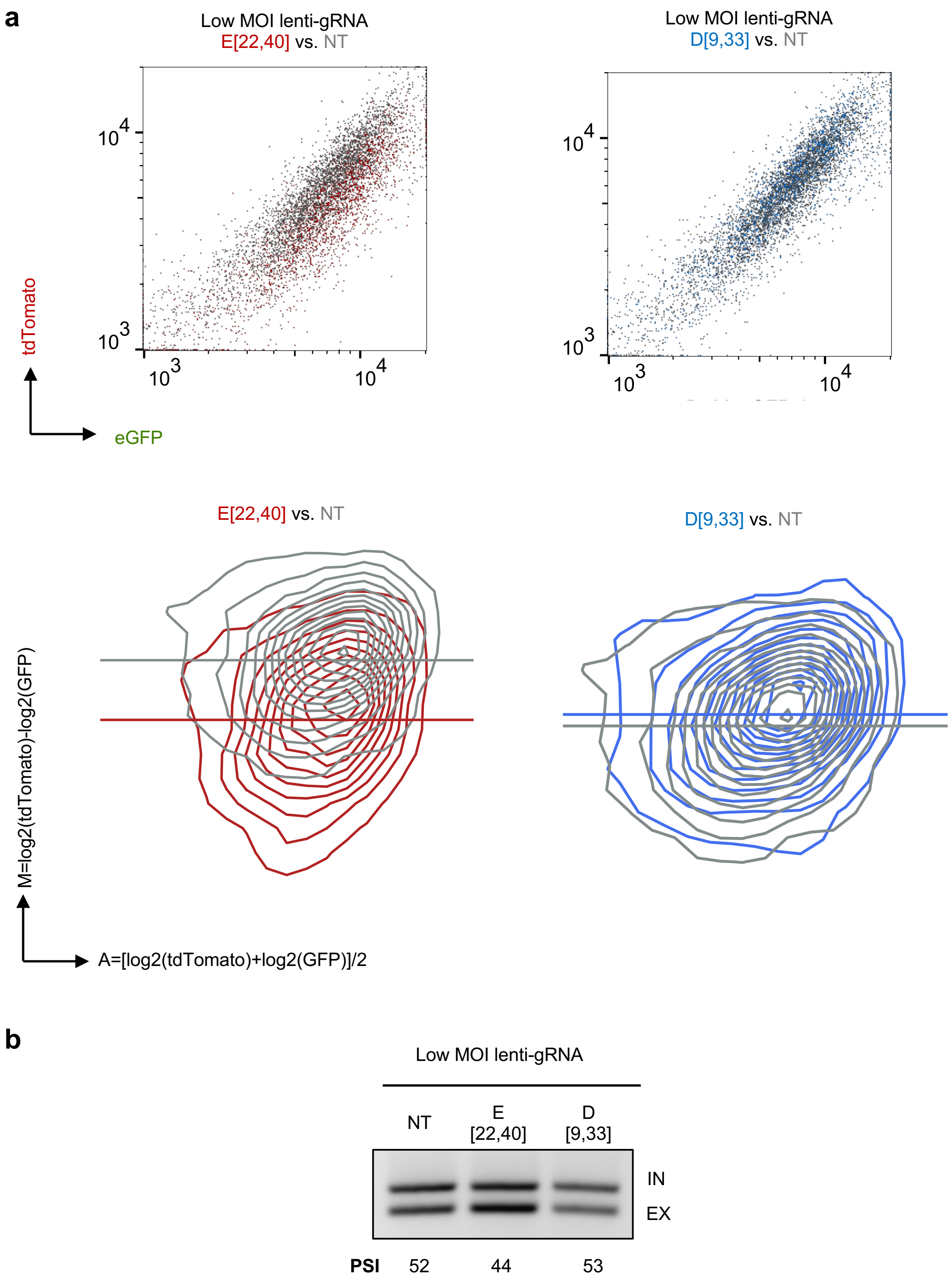


Supplementary Figure 9

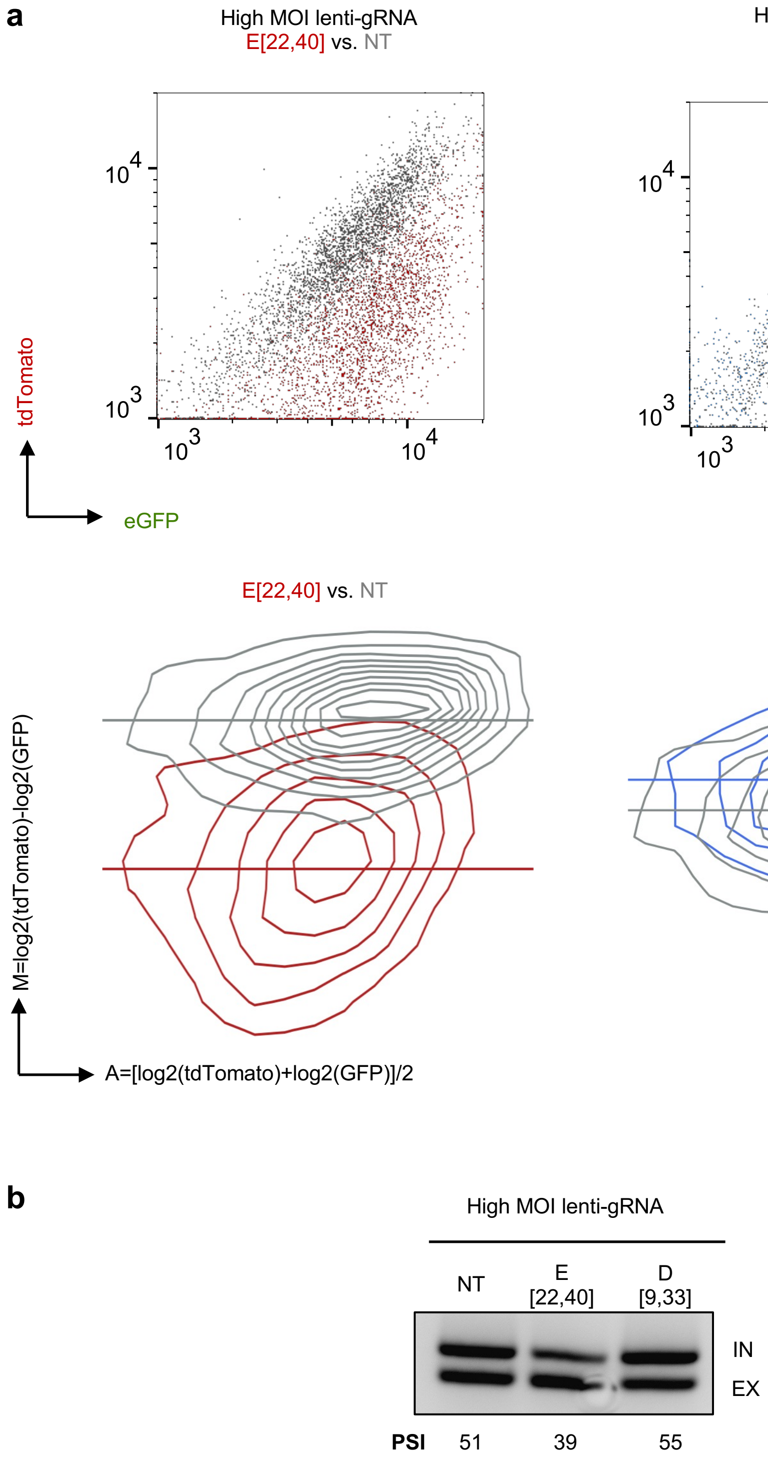


Supplementary Figure 10

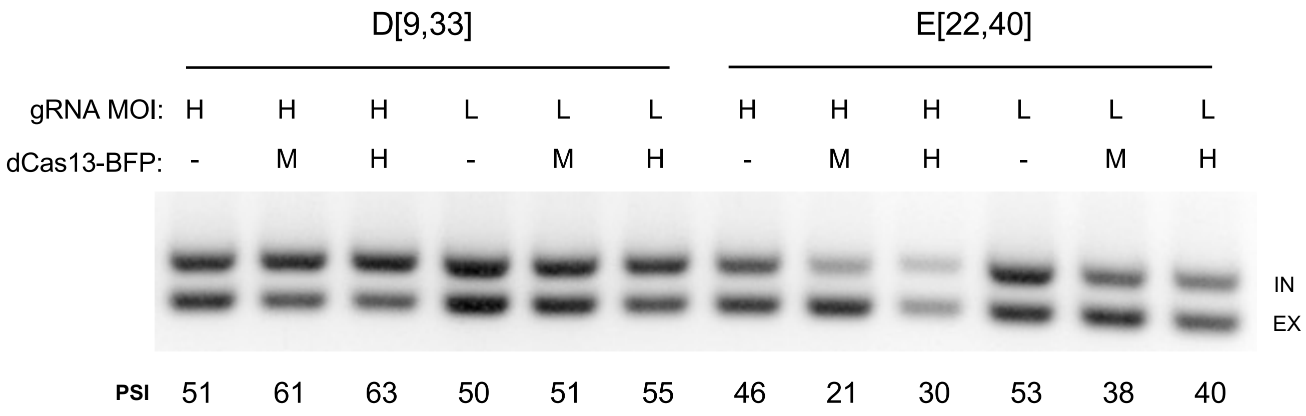


Supplementary Figure 11

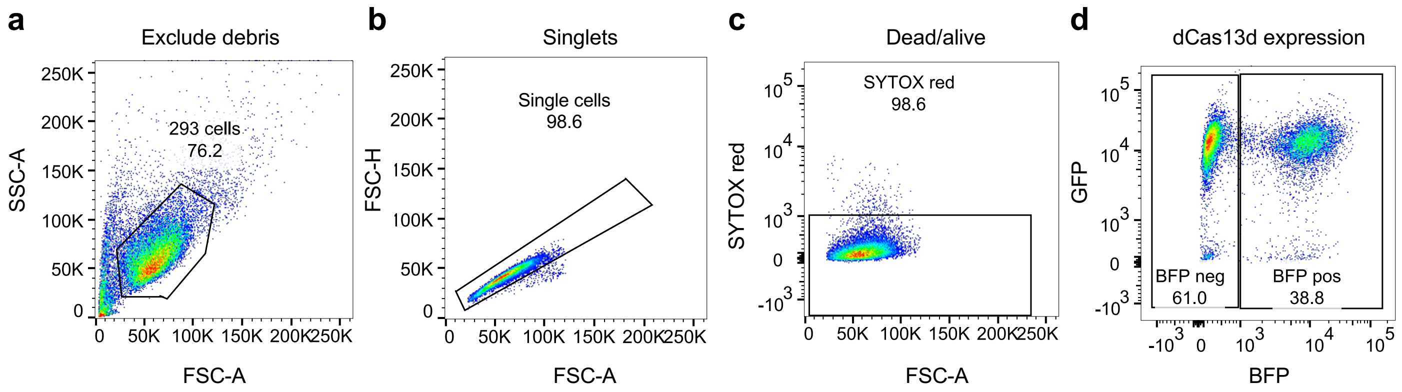


Supplementary Figure 12

**a**

| Dual-IN *SMN2* Samples | Total reads Replicate 1 | Perfect gRNA Reads Replicate 1 (%) | Total reads Replicate 2 | Perfect gRNA Reads Replicate 2 (%) |
|---|---|---|---|---|
| Unsorted | 9,832,395 | 7,726,408 (78.6%) | 13,099,961 | 10,729,246 (81.9%) |
| Top Bin BFP positive | 18,223,601 | 13,704,672 (75.2%) | 14,004,110 | 11,228,267 (80.2%) |
| Bottom Bin BFP positive | 14,074,967 | 11,298,635 (80.2%) | 10,388,855 | 8,768,692 (84.4%) |
| gRNAs Not Detected in Unsorted | - | 48 (2.5%) | - | 59 (3%) |

**b**

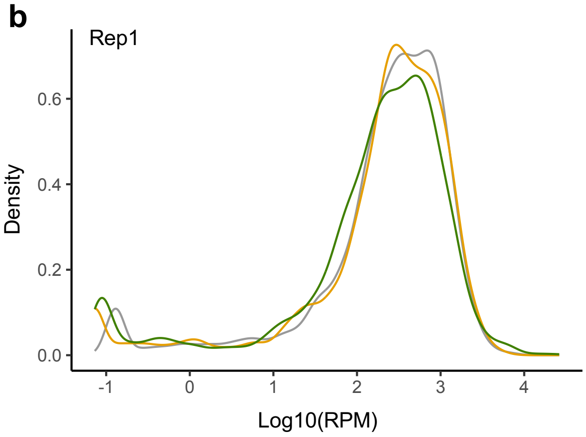


**c**

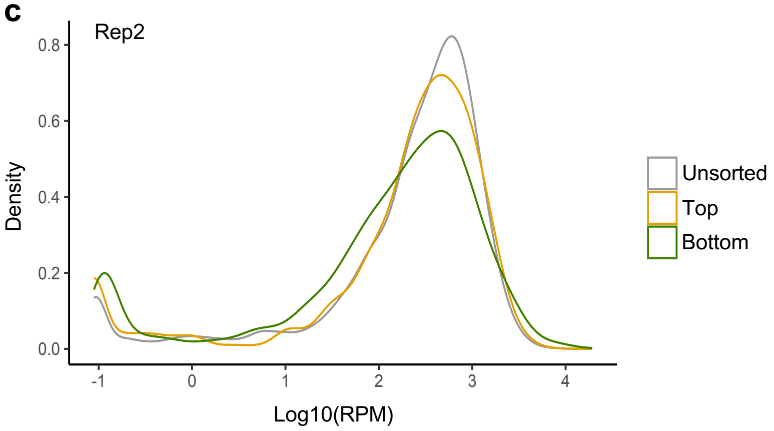


Supplementary Figure 13

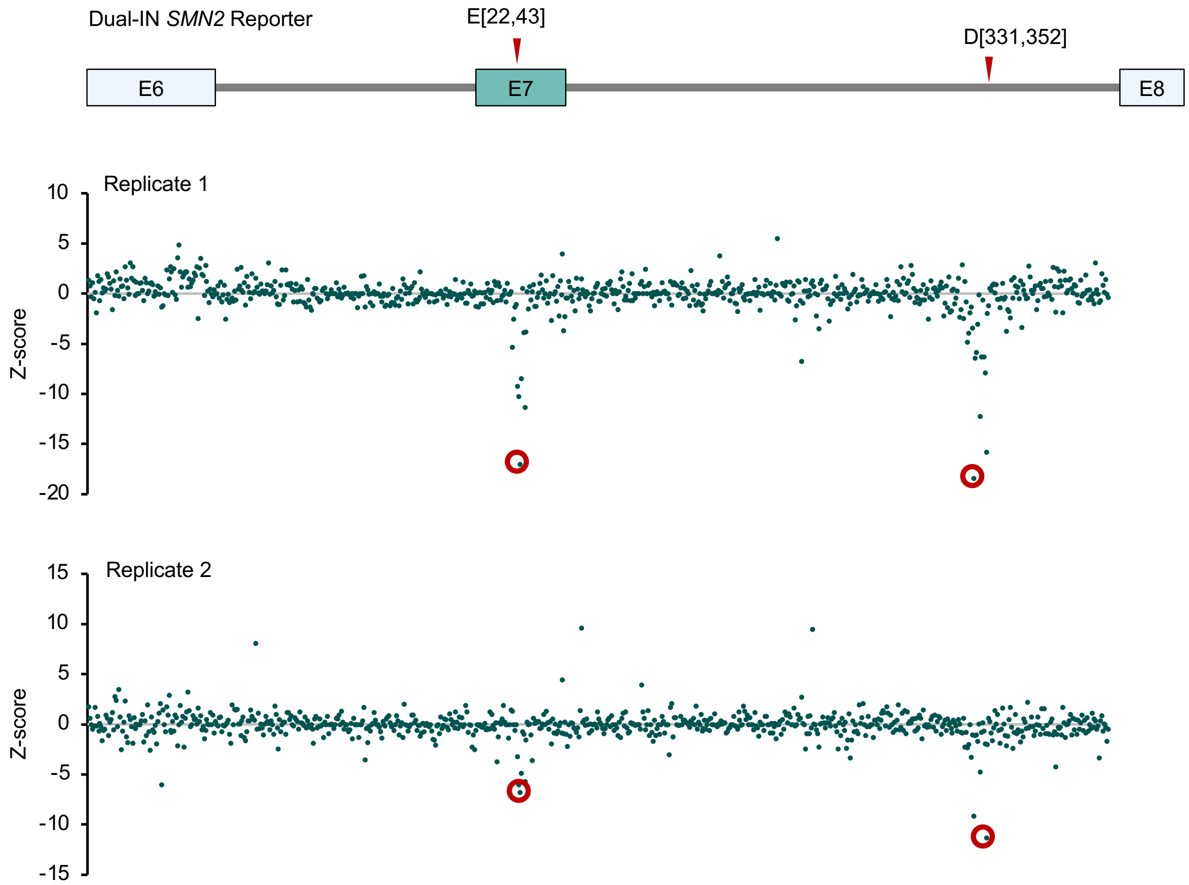


Supplementary Figure 14

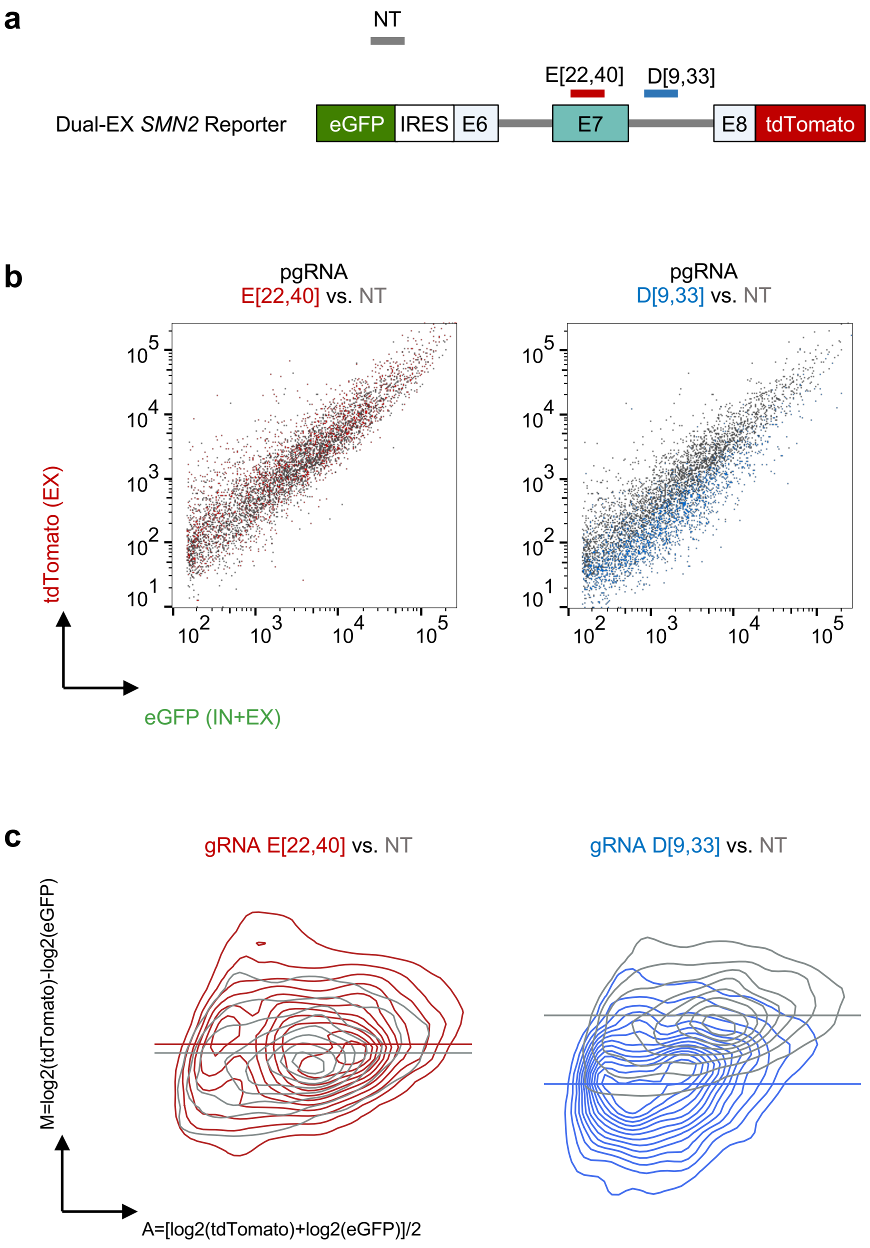


Supplementary Figure 15

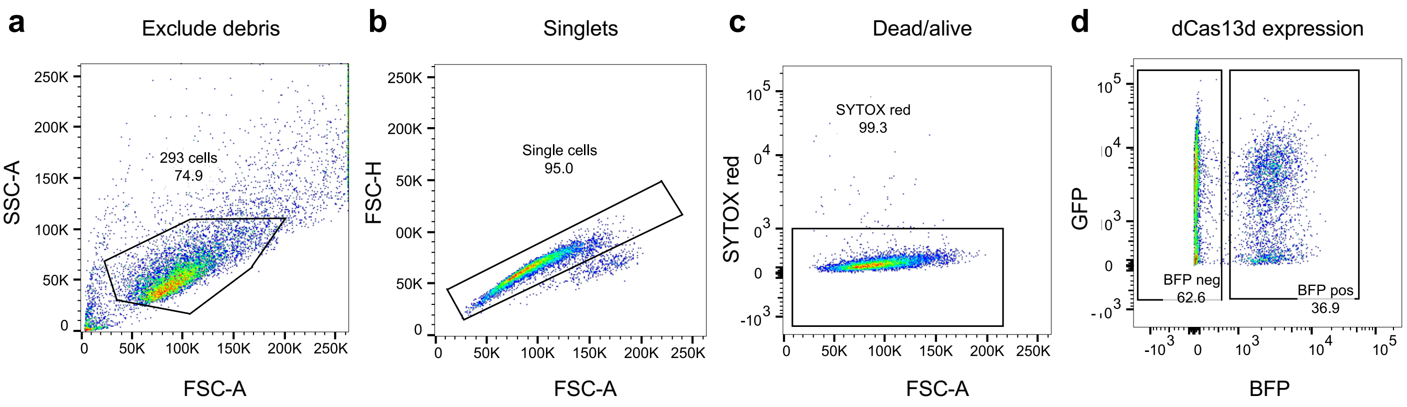


Supplementary Figure 16

**a**

| Dual-EX *SMN2* Samples | Total reads Replicate 1 | Perfect gRNA Reads Replicate 1 | Total reads Replicate 2 | Perfect gRNA Reads Replicate 2 |
|---|---|---|---|---|
| Unsorted | 11,229,347 | 8,962,706 (79.8%) | 13,978,926 | 10,797,580 (77.2%) |
| Top Bin BFP positive | 13,748,993 | 11,049,081 (80.4%) | 11,133,197 | 8,704,872 (78.2%) |
| Bottom Bin BFP positive | 13,369,246 | 10,645,588 (79.6%) | 15,697,524 | 12,139,831 (77.3%) |
| gRNAs not detected in unsorted | - | 46 (2.4%) | - | 54 (2.8%) |

**b**

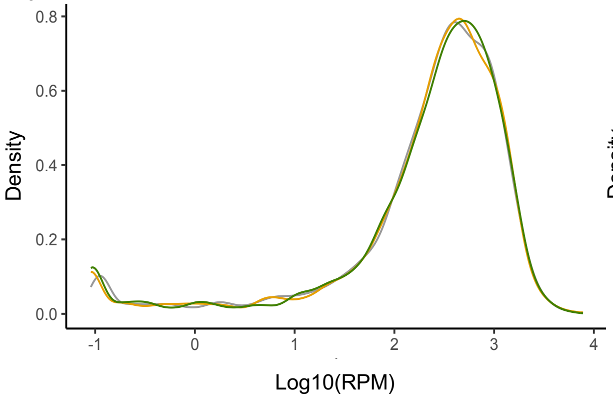


**c**

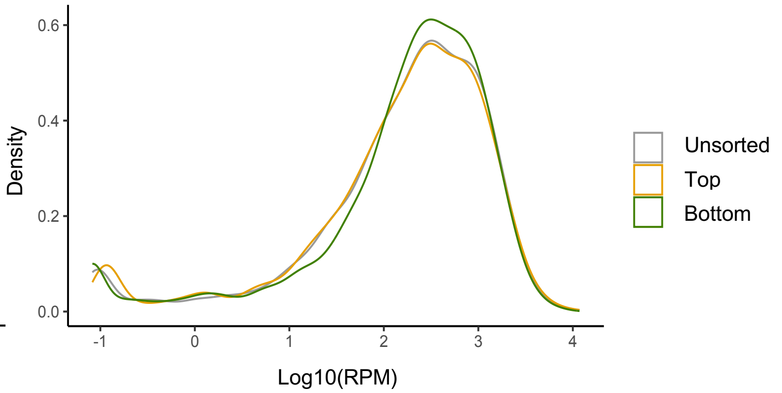


Supplementary Figure 17

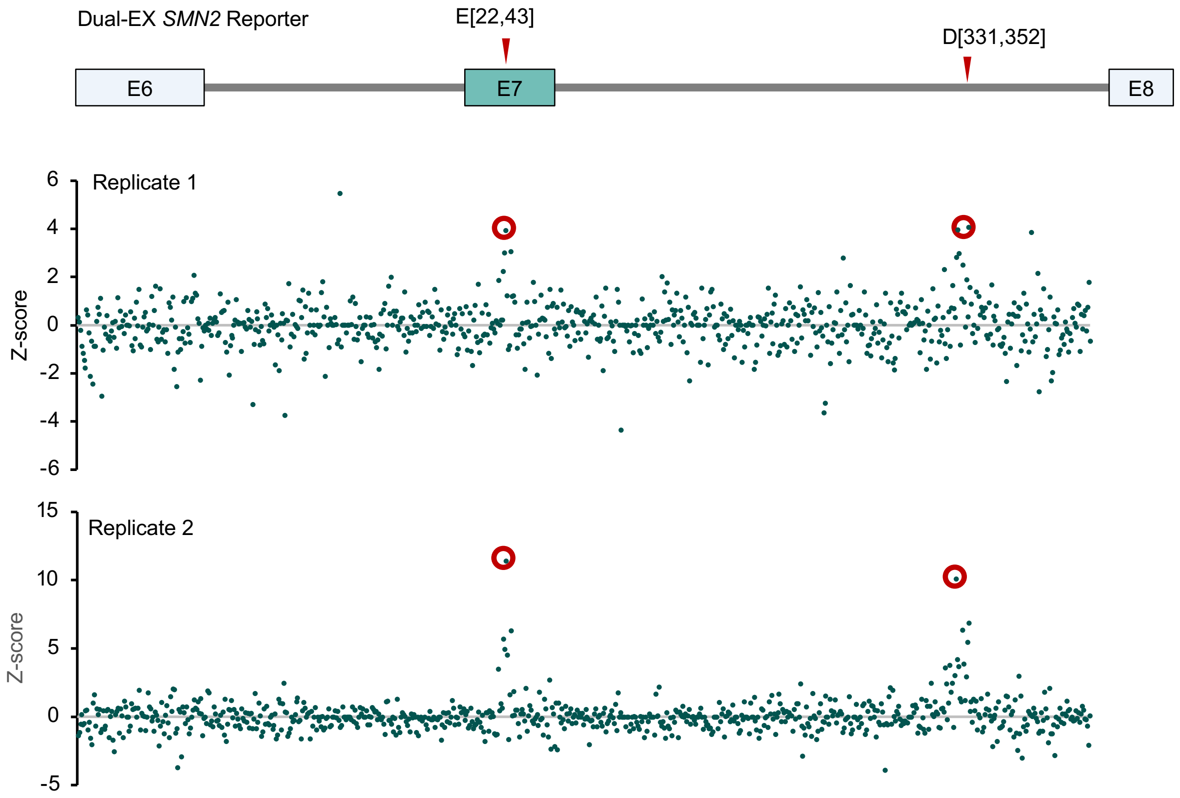


Supplementary Figure 18

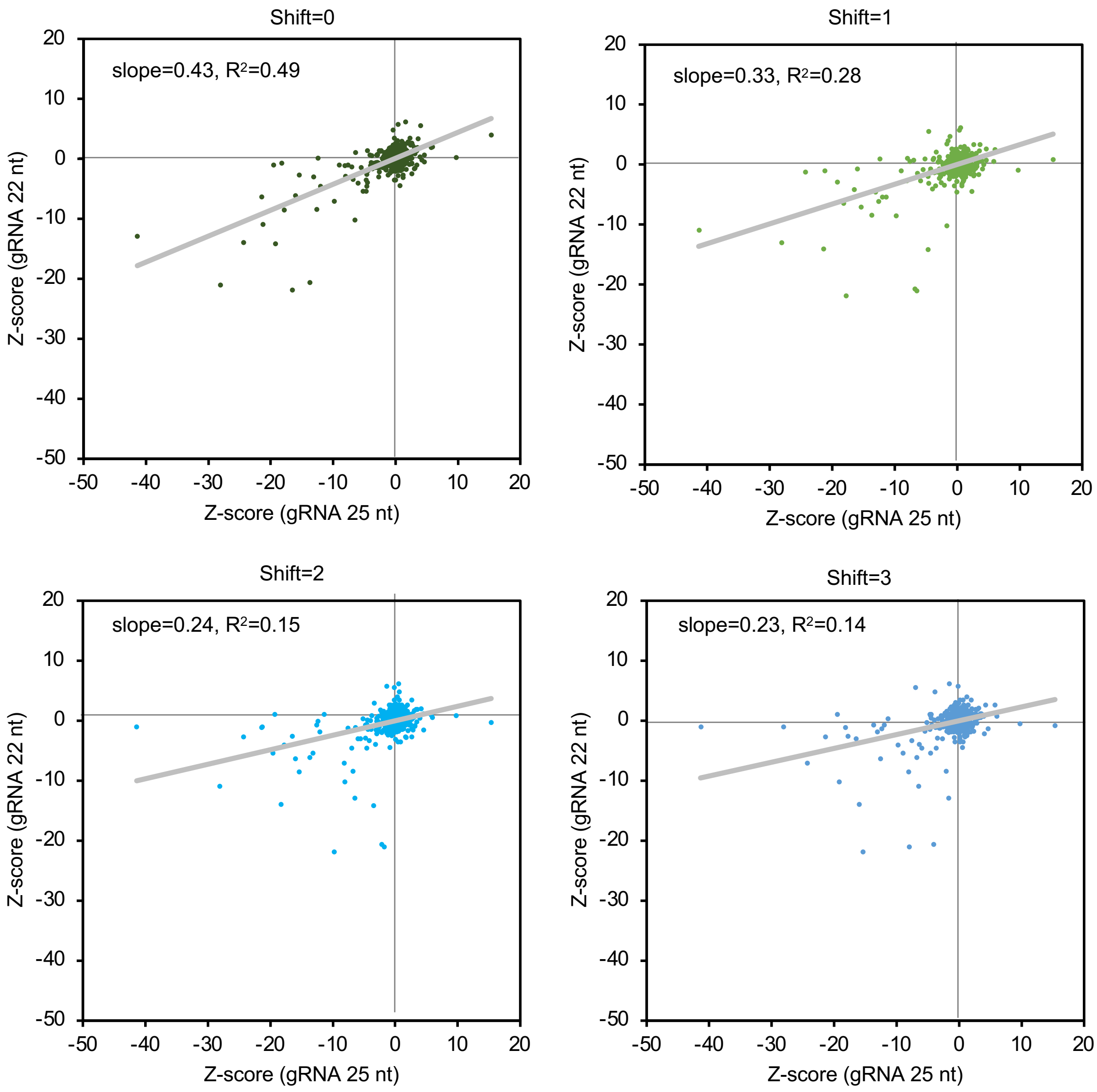


Supplementary Figure 19
